# Larger cerebral cortex is genetically correlated with greater frontal area and dorsal thickness

**DOI:** 10.1101/2022.05.19.492686

**Authors:** Carolina Makowski, Hao Wang, Anjali Srinivasan, Anna Qi, Yuqi Qiu, Dennis van der Meer, Oleksandr Frei, Jingjing Zou, Peter M. Visscher, Jian Yang, Chi-Hua Chen

**Author notes:** Correspondence to (C.H.C.).

## Abstract

Human cortical expansion has occurred non-uniformly across the brain. We assessed the genetic architecture of cortical global expansion and regionalization by comparing two sets of genome-wide association studies of 24 cortical regions with and without adjustment for global measures (i.e. total surface area, mean cortical thickness) using a genetically-informed parcellation in 32,488 adults. We found 393 and 756 significant loci with and without adjusting for globals, respectively, among which 8% and 45% loci were associated with more than one region. Results from analyses without adjustment for globals recounted loci associated with global measures. Genetic factors that contribute to total surface area of the cortex particularly expand anterior/frontal regions, whereas those contributing to thicker cortex predominantly increase dorsal/frontal-parietal thickness. Interactome-based analyses revealed significant overlap of global and regional genetic modules, enriched for neurodevelopmental and immune system pathways. Consideration of global measures is important in understanding the genetic variants underlying cortical morphology.

---

The human cerebral cortex has undergone an extraordinary expansion compared to other mammalian species, mirroring the development of many complex traits unique to modern-day humans. The cerebral cortex is a layered, folded sheet of gray matter and its size can be measured by surface area tangentially and thickness radially, with differential neurodevelopment programs shaping and regulating these two cortical measures^1^. The product of these two measures roughly corresponds to measures of cortical volume^2^. Surface area expansion in humans, particularly in functionally unique association areas, has undergone a 2000-fold increase relative to mice, compared to an only 3-fold increase in cortical thickness^3^. The prefrontal cortex in particular may have disproportionately expanded in humans compared to non-human primates^4^. Motivated by the human cortical expansion observed across species, here we study phenotypic variation of the cortex in humans. Differentiating between the genetic variants linked to regionalization and global size of the cortex in humans may give us insights into the functional specialization required in development to define unique brain regions that are important to our understanding of cognition and brain disorders.

It is known that larger mutations within the genome, such as copy number variants (CNVs), have quite profound effects on both global and regional brain morphology, and contribute to several neurodevelopmental conditions^5^. Twin-based heritability estimates have suggested genetic overlap between regional surface area/thickness and global measures^6^. However, it is unknown which genetic variants contribute to this overlap and how these genetic variants shape global and regional brain development. A study by Shin and colleagues began to address this question of the different information that can be gleaned from the genetic architecture of global compared to regional features of the cortex, where they carried out a GWAS in 23,000 individuals on the top two principal components of cortical surface area data^7^. As expected, the first principal component largely captured global features of the cortex, whereas the second principal component captured occipital/visual cortex-specific surface area. Their study found that the genetic architecture of these two components differentially associated with complex traits, such that the first “global” component seemed to map onto the genetic architecture of general cognitive and learning ability, whereas the second “regional” occipital component had higher genetic correlations with psychiatric disorders. It remains to be seen how the genetic architecture of global surface area compares and contrasts with other regions of the brain, particularly newer cortical regions subserving higher cognitive function. Additionally, cortical thickness also shows unique regionalization patterns which shape the functional boundaries of the brain^8^.

In this study we assessed the relationship between global measures (e.g. total surface area and mean cortical thickness across the brain) and regional patterns of genetically-informed brain morphology, where brain regions were defined by hierarchical clustering of twin data as presented by our group previously^9–11^. The genetically-informed brain parcellations applied to this study adhere to known genetic patterning of the cortex, including anterior-posterior and dorsal-ventral developmental axes of surface area and cortical thickness, respectively. To do so, we compared genetic variants and genes associated with 12 surface area and 12 cortical thickness regions from two sets of GWAS, with and without adjustment for global brain measures. Global adjustment is typically done in brain imaging studies to account for the fact that some individuals will have larger morphological features simply due to having a larger brain. We employed an interactome-based gene mapping approach to investigate the genetic overlap or separation in genes associated with global measures and different brain regions. We expect that global measures will have higher genetic overlap with association areas (e.g. prefrontal cortex) compared to primary sensory cortices (e.g. occipital cortex), given the disproportionate expansion of association regions that contribute to the protracted course of human brain development, and in turn, higher cognitive function.

## RESULTS

### Sample

Our final sample included brain imaging and genetic data from 32,488 individuals (mean age=64.2 [range: 45.1-81.8, SD: 7.5], %female=52.2) from the UK Biobank. See Supplementary Table 1 for demographics and descriptives of the regional brain data.

### GWAS analyses and workflow

Twelve regions of interest were extracted per hemisphere based on two genetically-informed atlases for cortical thickness and surface area for both discovery and replication cohorts. These atlases have been previously developed by our group^9,10^, using a data-driven fuzzy clustering technique to identify parcels of the human cortex that are maximally genetically correlated based on the MRI scans of over 400 twins.

All GWAS analyses were carried out on pre-residualized brain phenotypes, adjusted for age, sex, scanner site, a proxy of scan quality (FreeSurfer’s Euler number)^12^, presence of a brain disorder, and the first ten genetic principal components. See Online Methods for more details. We denote one set of analyses as GWAS*r* for regional associations, which also adjusts for global measures (i.e. total surface area, mean whole brain thickness) in the pre-residualized regional brain phenotypes, and the second set of regional associations as GWAS*g+r*, which *includes* global measures (i.e. does not adjust for globals). We also include GWAS of global measures (i.e. total cortical surface area, mean whole brain cortical thickness), which we denote as GWAS*g*. Thus, in total we analyzed 50 GWAS (24 GWAS*g+r*, 24 GWAS*r*, 2 GWAS*g*). Results from GWAS*r* and GWAS*g* have been previously reported by our group^11^. In this study, we include only UKB participants with European ancestry, although our previous work has also shown generalizability of GWAS*r* results to a smaller sample of individuals with more diverse ancestry and generalization to childhood/early adolescence^11^. See Figure 1A for a visualization of these different GWAS analyses.

**Figure 1:**
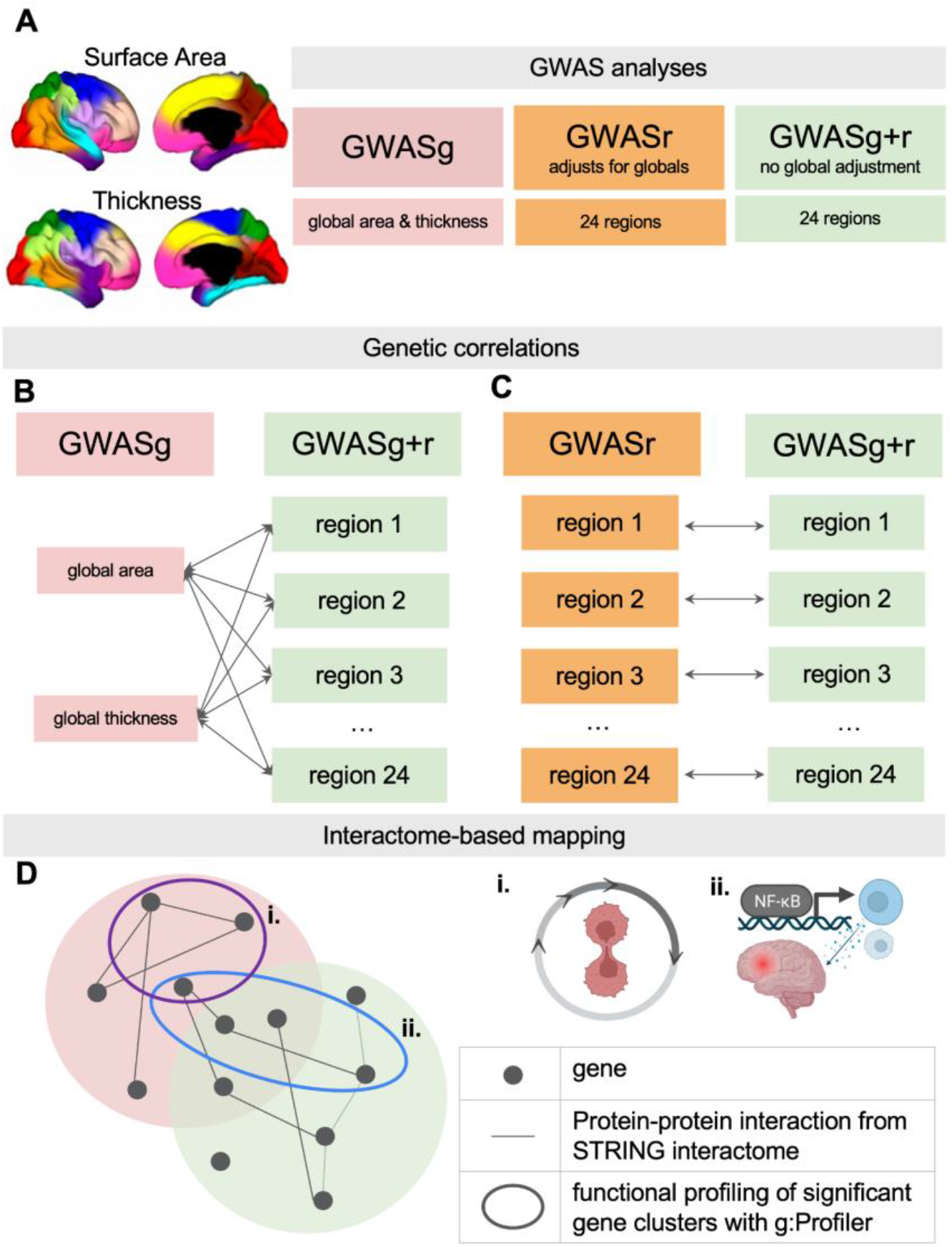
Methods workflow. *Panel A*: Three sets of GWAS on GWAS of global measures (GWASg), regional GWAS adjusting for global measures (GWASr) and regional GWAS not adjusting for global measures (GWASg+r). *Panels B and C*: Two sets of genetic correlation analyses with LDSC. *Panel D*: Interactome-based mapping of gene lists comparing GWASg and GWASg+r phenotypes. For select phenotype pairs, functional profiling was then carried out with g:Profiler for functional interpretation of meaningful gene clusters within the Venn Diagram, for example i. cell cycle-related annotations; ii. inflammatory pathways, such as processes involving Nuclear Factor Kappa B (NF-κB).

### Global measures are significantly associated with regional phenotypes

We first estimated the effects of the two global measures and chosen covariates (age, sex, scanner, brain diagnosis, Euler number, 10 PCs) on our cortical phenotype data using four sets of tests: the Akaike and Bayesian Information Criterion (AIC, BIC), ANOVA F tests, and Least Absolute Shrinkage and Selection Operator (LASSO). See Online Methods for more details on these tests. AIC and BIC values were both smaller across all phenotypes in the full model compared to the reduced model which excluded global measures, suggesting that the full model including global surface area/thickness provided a significantly better fit to the cortical phenotype data. ANOVA F-tests provided a similar conclusion, where the null hypothesis that the simpler (reduced) model is as good as the full model was rejected for all phenotype models. Finally, LASSO consistently selected the global measures as an important contributor to our models for each cortical phenotype. We also ran LASSO easing restrictions and allowing the operator to select from all other 15 covariates in the model. This approach showed that age, sex, and the Euler number (a proxy of image quality) were important contributors to our models for nearly all cortical phenotypes, and scanner contributed notably to about half of the phenotypes. See Supplementary Table 2.

### GWAS of cortical phenotypes for both GWAS*r* and GWAS*g+r*

We compared the number of significant loci associated with our cortical phenotypes for GWAS*r* (Supplementary Tables 3, 4) and GWAS*g+r* (Supplementary Tables 3, 5). In our latest work on the same sample^11^, we reported 393 significant loci associated with our 24 cortical phenotypes, after clumping with PLINK^13^ (r^2^=0.1, distance=250kb) and thresholding at p<5e-8, of which 361 loci had unique rsIDs (i.e. 8% duplicated loci). Further, for GWAS*g* we found 27 and 20 variants significantly associated with total cortical surface area and mean cortical thickness, respectively. For GWAS*g+r*, we obtained 756 significant loci across the 24 cortical phenotypes, of which 419 were unique; in other words, 45% were duplicated loci, due to the recounting of loci underlying global measures. Across the unique genome-wide significant loci for GWAS*r* and GWAS*g+r*, 267 and 225 were found to be LD-independent by clumping all phenotypes together in PLINK^13^ (r^2^=0.1, distance=250kb). Miami plots comparing these two GWAS approaches, as well as number of hits per region, can be found in Figure 2A and 2B for area and thickness, respectively. Gene ontology (GO) analysis results for both GWAS*r* and GWAS*g+r* can be found in Figure 3A and 3B, respectively (Supplementary Tables 6, 7). SNP-based heritability range estimates from LDSC are as follows and can be found in Supplementary Table 8: GWAS*r*, area from 0.23 to 0.36; GWAS*r*, thickness from 0.15 to 0.25; GWAS*g+r*, area from 0.35 to 0.39; GWAS*g+r*, thickness from 0.21 to 0.31. Consistent with previous work, after regressing out globals, heritability estimates drop on average by 0.09 and 0.07 for area and thickness respectively (compared to ∼0.2 using twin heritability estimates6).

**Figure 2.**
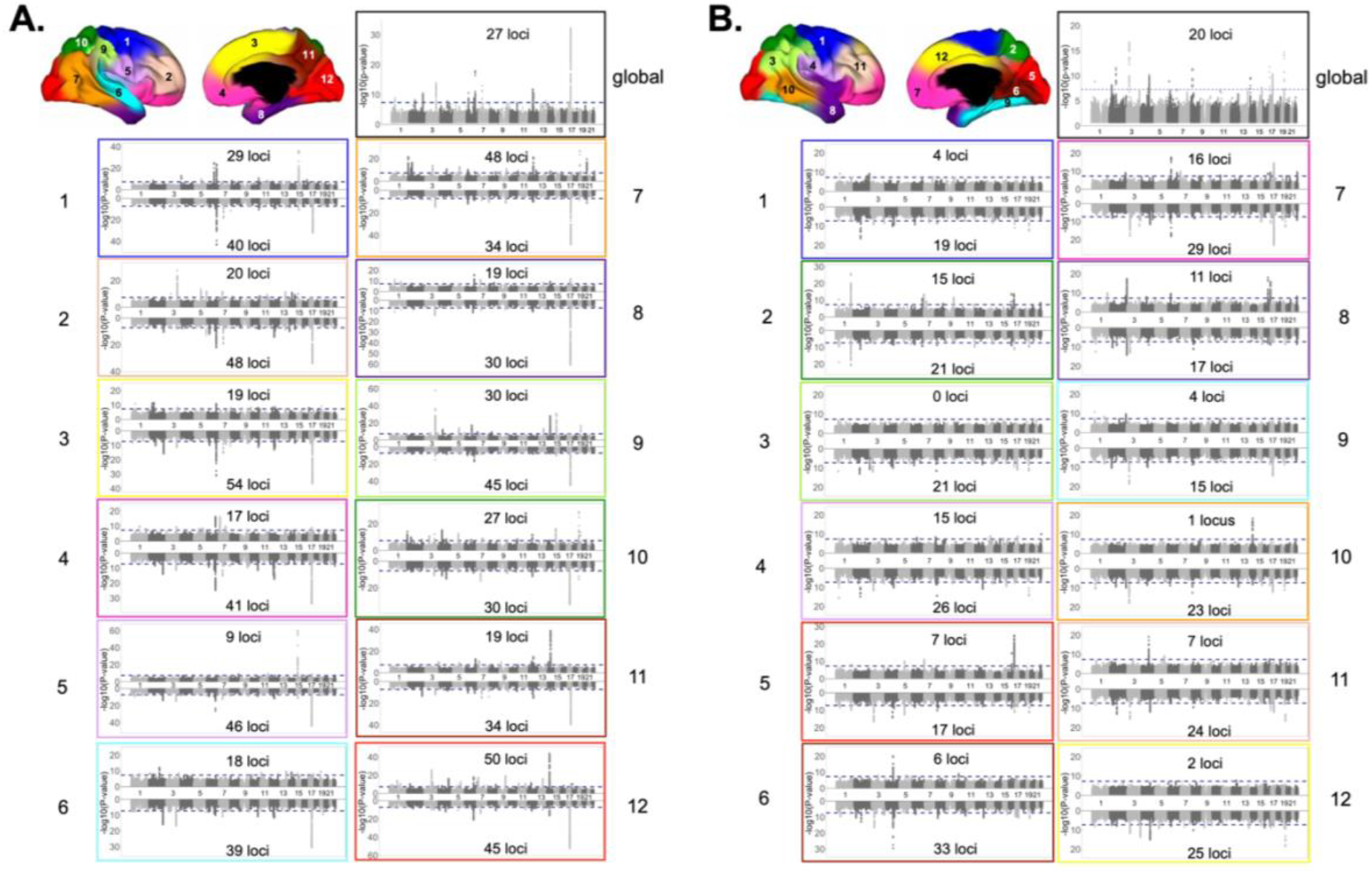
Miami plots colored by atlas region for *Panel A*. surface area and *Panel B*. cortical thickness. Top half of Miami plot for GWAS*r*, and bottom half for GWAS*g+r*. Manhattan plots for global surface area and mean cortical thickness are included in the top right of each panel. Number of significant loci (defined by plink, r^2^=0.1, 250kb) for each analysis are included in each subplot. Numeric and colored labels on brain maps correspond to the same number/color of each subplot.

**Figure 3.**
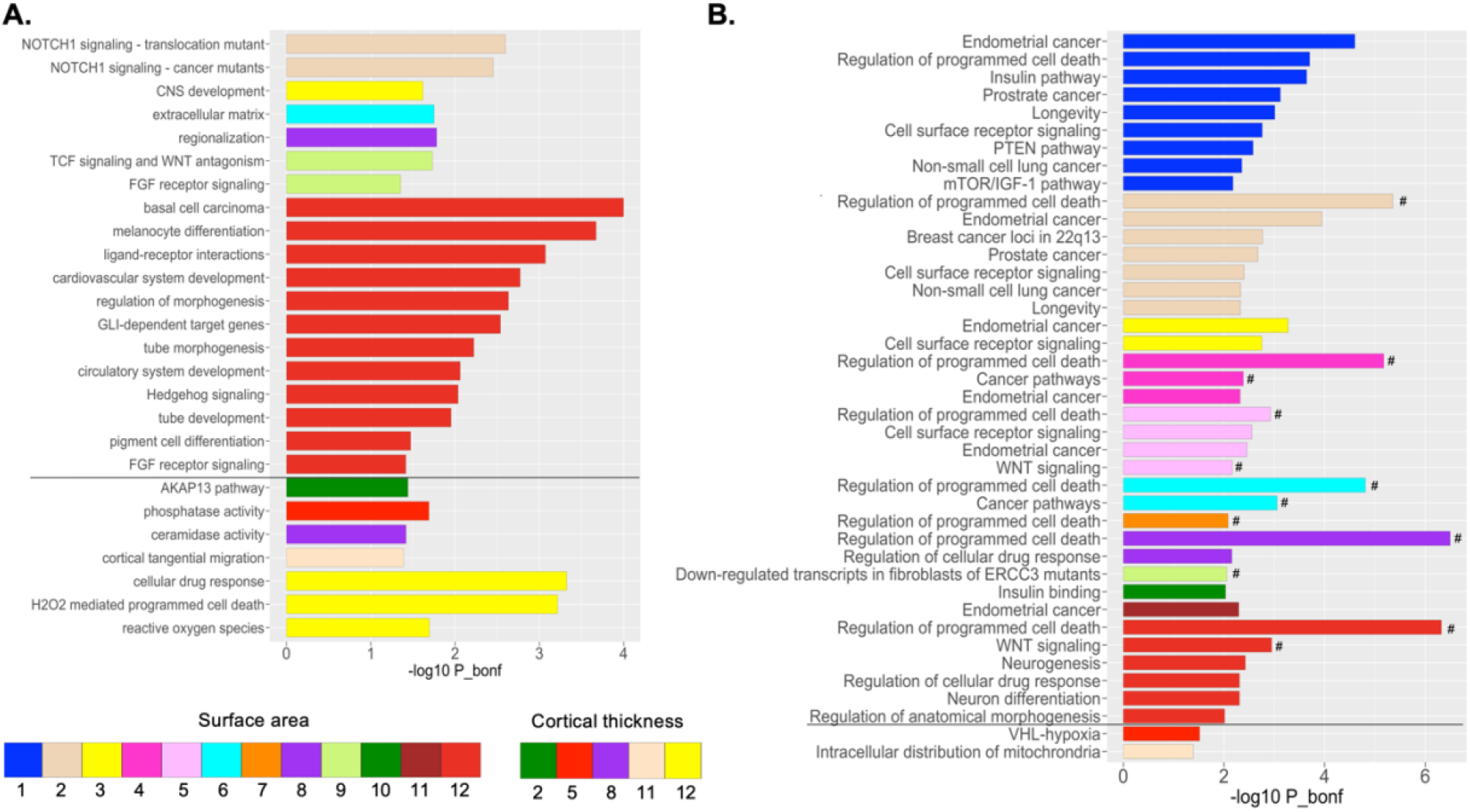
Bonferroni-corrected GO terms (p<0.05, unless otherwise specified) for *Panel A*. GWAS*g* (adapted from ^11^) and *Panel B*. GWAS*g+r*. Bars are color-coded by cortical region, where the region numbers included in the legend correspond to region numbers in Figure 2. Results above the horizontal black line represent surface area regions, and regions below for cortical thickness. Hashtags (#) in *Panel B* reflect terms that are also significantly associated with global surface area. For readability, only terms with Bonferroni-corrected p<0.01 are displayed in *Panel B*. See full list of Bonferroni-corrected terms (p<0.05) in Supplementary Table 7.

As has been shown previously^7,14^, we observed large effects in global surface area for loci within the 17q21.31 inversion region. This signature on chromosome 17 was seen across all 12 surface area phenotypes for GWAS*g+r* analyses but not for GWAS*r*, emphasizing the additional signal that global brain measures offer in detecting genetic loci shaping the brain. The most significant loci associated with any particular region also varied by approach; for GWAS*r*, the strongest signal was detected for rs7182018 (p=2.2e-60, variance explained 0.83%) associated with pars opercularis area within 15q14; for GWAS*g+r*, rs55751924 (p=1.33e-61, variance explained 0.84%) associated with anteromedial temporal area on 17q21.31; and for GWAS*g*, rs593720 (p=4.94e-33, variance explained 0.44%) associated with total surface area also within 17q21.31. Intriguingly, we did not see a strong signal on 15q14 associated with global area, despite its contribution to the surface area of pars opercularis of the inferior frontal cortex. For mean cortical thickness, the strongest association was found with rs694611 (p=2.10e-17, variance explained 0.22%) on chromosome 3, a genetic locus that was also found to be significantly associated with the thickness of most phenotypes in GWAS*g+r*. This SNP lies within the *RPSA* gene, which has recently been shown to have an important role in early cortical development, particularly for dendritic spine morphology and cortical layering^15^.

### Comparing genetic architectures to quantitatively assess the relationship between global and regional measures

*Comparing GWASg with GWASg+r*. We calculated LD score regression (LDSC)-based genetic correlations (rg) between GWAS of each global measure (GWAS*g*: total surface area and mean thickness) and GWAS*g+r* results for the 24 cortical phenotypes to better understand the association of global measures with regional brain morphology (Figure 1B). As can be seen in Figure 4A, the highest genetic correlations with global area were found with anterior/frontal regions; and for global thickness, highest correlations were with dorsal regions, specifically fronto-parietal (rg ranges for i. global area with regional area: 0.80 to 0.93; ii. global area with regional thickness: -0.20 to -0.55; iii. global thickness with regional area: -0.26 to -0.36; and iv. global thickness with regional thickness 0.71 to 0.92; all p’s<4e-4) (Figure 3A, Supplementary Table 9).

**Figure 4.**
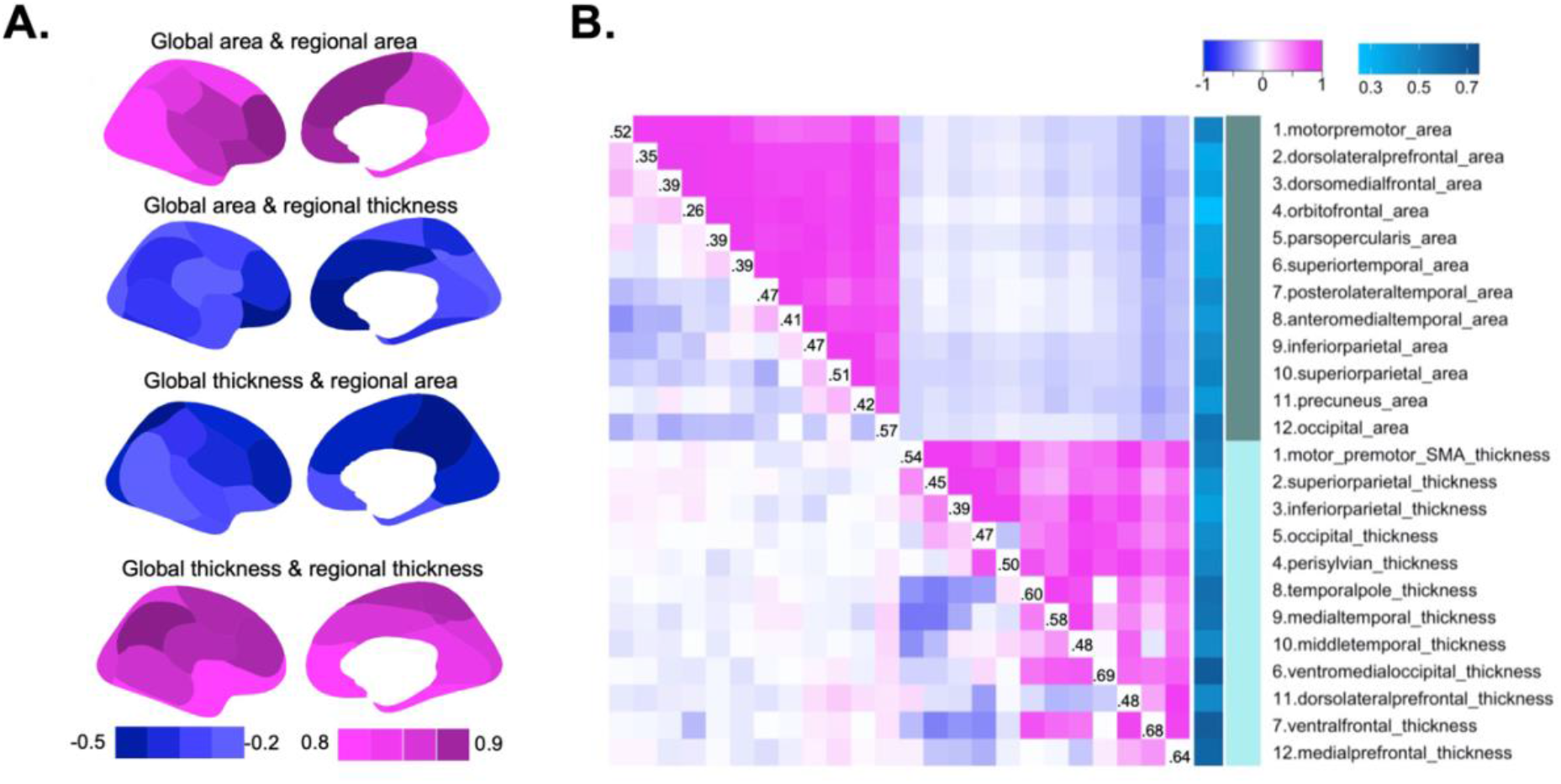
Phenotypic and genotypic relationships. *Panel A*: Genetic correlations between GWAS summary statistics of global measures and regional GWAS*g+r. Panel B*: Lower triangle of matrix displays phenotypic correlations when globals are regressed out, whereas upper triangle shows phenotypic correlations when globals are *not* regressed out during pre-residualization. The diagonal labels reflect genetic correlations between GWAS*r* and GWAS*g+r* results, which are also depicted in the blue color bar to the right of the matrix.

#### Comparing GWASr and GWASg+r results

We also used LDSC to compute genetic correlations between GWAS*r* and GWAS*g+r* results for each of the 24 brain regions as another way of understanding and visualizing the genetic contributions of global measures to regional brain morphology. Genetic correlation results can be found in Figure 4B and Supplementary Table 8, where we find genetic correlations between GWASr and GWASg+r of the same phenotypes range from 0.26 to 0.57 for surface area, and 0.39 to 0.69 for cortical thickness (mean rg: 0.49, mean standard error: 0.042, all p’s<4e-7). Generally, there were higher genetic correlations for cortical thickness compared to surface area, and higher associations were found for primary sensory/motor regions compared to association and frontal regions. This approach gave expected results, with a reversal of patterns shown in the section above. For example, of the area phenotypes, occipital area has the highest rg between GWASr and GWASg+r but lowest rg between GWAS*g* and GWAS*g+r*, suggesting that occipital area is least affected by adjustment for total surface area. In other words, genetic factors that influence occipital area are less similar than those that influence total surface area.

For completion we also looked at the phenotypic correlations between all 24 regions comparing GWAS*r* and GWAS*g+r*, which can be found in Figure 4B. We have previously published the phenotypic correlations for GWAS*r*^*10,11,16*^. Notably, much more uniform phenotypic correlation structures emerge when globals are not regressed out, likely due to the shared global genetic variants underlying many regions. Specifically, we observe largely positive correlation structures within a trait (e.g. one area region with another area region), but negative correlations across traits (e.g. one area region with a thickness region) (see Figure 4B).

### Overlap in genetic modules between global and regional brain measures

We carried out a network separation analysis adapted from Menche and colleagues^17^ to see whether the genes underlying global measures (i.e. from GWAS*g*) were overlapping or separated from genes underlying regional brain measures (i.e. from GWAS*g+r*). We assessed the degree of overlap by calculating separation (SAB) and mean shortest distance (dAB) between each global (A) and regional brain region (B) pair. Calculations for these metrics are outlined in Online Methods.

We used MAGMA to define genes included in the genetic modules for global and regional phenotypes. SNPs were selected that were within exonic, intronic and untranslated regions as well as SNPs within 50 kb upstream and downstream of the gene, a window size that has been used in previous cortical GWAS^14^. A summary of associated MAGMA genes are included in Supplementary Table 10, and lists of genes and statistics are found in Supplementary Tables 11 and 12 for GWASr, GWAS*g*, and GWAS*g+r*. Consistent with the genetic correlation patterns, genetic modules between global and regional phenotypes, for both area and thickness, showed significant overlap, as demonstrated by negative separation values (sAB z-scores, area range: -3.84 to -14.70; thickness range: -4.13 to -14.51; all p’s<2e-5). See Supplementary Table 13 for separation and shortest distance statistical results. Overlap in genetic modules was particularly high between global area and area of prefrontal regions (e.g. orbitofrontal, pars opercularis), as well as between global thickness and thickness of fronto-parietal regions (e.g. dorsolateral prefrontal, inferior parietal), consistent with genetic correlation results presented in Figure 4A. Genes that were mapped by MAGMA but not found in the STRING interactome database are excluded and listed in Supplementary Table 14; the function of many of these excluded genes remains largely unknown as they are pseudogenes or non-coding RNA genes. The genetic overlap between global and regional measures are displayed for two exemplary pairs in Figure 5: i) global surface area and dorsolateral prefrontal area, and ii) global thickness and dorsolateral prefrontal thickness. Functional profiling of gene lists from each separate module (e.g. brain region) and overlapping genes was completed with g:Profiler^18^, and revealed many overlapping genes were important in immune function and early neurodevelopmental processes. Network figures were created using Cytoscape^19^ (Figure 1D).

**Figure 5.**
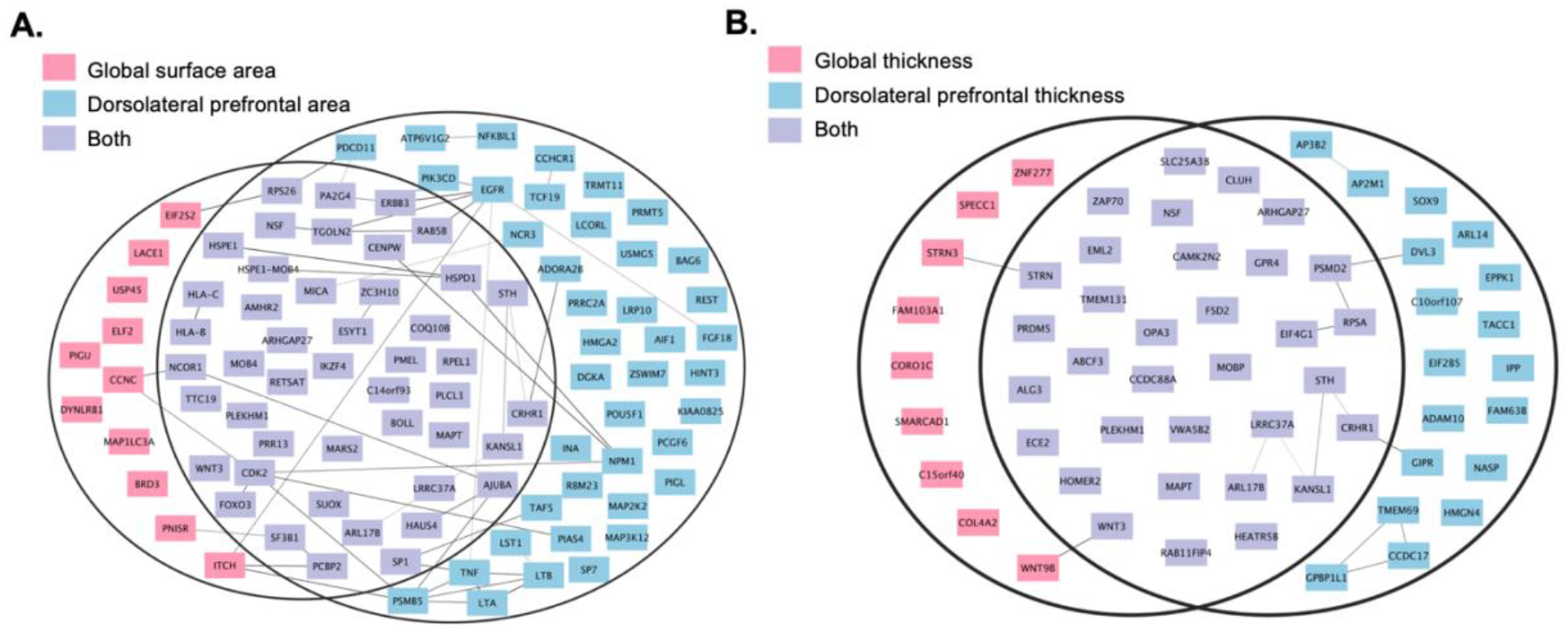
Venn Diagrams of overlap in genetic modules for two exemplary pairs. *Panel* A: global surface area and dorsolateral prefrontal cortical area, and *Panel B:* global thickness and dorsolateral prefrontal thickness. Nodes in the graph represent genes, and edges (grey lines) represent connections between genes as defined by the STRING interactome.

In addition to using gene lists generated by MAGMA, we also computed separation statistics using genes mapped by FUMA, which includes intergenic mapping (Supplementary Tables 13, 16-18). Results and interpretation from FUMA gene mapping were similar to those obtained with MAGMA.

### Finding common genetic loci between the two sets of analyses

In addition to assessing overlap in genetic modules between traits with an interactome-based mapping approach, we also compared the list of significant loci (clumped with PLINK^13^ with r^2^=0.1, distance=250kb, p<5e-8) between GWAS*r* and GWAS*g+r*. We first identified common SNPs based on shared rsID by region, which identified 31 SNPs. Given that significant SNPs in LD with the identified SNPs may also be shared between analyses, we also proceeded to use conditional and joint (COJO) analysis^20^ of GWAS summary statistics implemented in GCTA to map shared SNPs for each cortical phenotype. We defined SNPs as being shared between the two approaches as those that were no longer genome-wide significant (i.e. p>5e-8) in GWAS*g+r* when conditioned on the significant loci defined by the GWAS*r* analysis of region *k*. There were 70 SNPs identified through this approach. These SNPs are listed in Supplementary Table 15.

### Sensitivity analysis using modified global measures

To ensure our genetic correlation estimates were not biased by the inclusion of a given region of interest in our global measure, we estimated the genetic correlation between GWAS*g+r* for dorsolateral prefrontal area and GWAS*g* for a modified global surface area measure where we excluded dorsolateral prefrontal area from the calculation. Results were highly similar (rg between dorsolateral prefrontal and original global measure: 0.93 [s.e.: 0.013]; rg between dorsolateral prefrontal and modified global measure excluding dorsolateral prefrontal area: 0.91 [s.e.: 0.0093]).

## DISCUSSION

This work shows the importance of considering global cortical measures in understanding the genetic variants underlying cortical morphology. Total surface area and mean cortical thickness are important contributors to phenotypic variation across all 24 brain phenotypes as identified by various statistical model fits, including LASSO. We show that human cortical area expansion is most associated with anterior/frontal expansion, and thicker cortex is most associated with dorsal/fronto-parietal thickening across individuals. Lowest genetic overlap was found between total surface area and occipital area, concordant with previous reports using transcriptome data in humans^21,22^ and non-human primates^23^ to show that the genetic patterning of the visual cortex is most distinct from other cortical regions. Given the higher genetic overlap we found between global measures and association brain areas (e.g. fronto-parietal regions), we investigated this further with a database of protein-protein interactions to understand the biological pathways that may contribute to the unique expansion of brain regions subserving higher cognitive function. Our interactome-based approach highlights the important information that can be gleaned from looking at the interactions between gene products rather than individual gene lists to assess shared genetic architecture and in turn, shared biological mechanisms, between two traits. Further, mapping such gene networks and their interactions can help us pinpoint specific functional modules that are important in understanding and treating neuropsychiatric disorders^24–26^.

Our work highlights both pros and cons for the use of global measures in GWAS models. The benefits of adjusting for global brain measures allow us to identify variants associated with regional measures relative to global measures. On the other hand, adjusting for global measures may reduce signals of interest, especially for regions such as the prefrontal cortex that are highly genetically correlated with global measures. Indeed, the strong impact of global brain measures on regional brain results has been demonstrated in association with many copy number variants^5^, which some of our common variants may also tag. Adjustment for global measures may also introduce a potential collider bias in the case that global measures along with genetic variants directly affect regional morphology; exploration of more complex statistical models to capture the potential bi-directional relationship between regional and global cortical morphology is warranted in future studies.

Many of the overlapping genes that were linked to both global and regional brain measures seem to be important in regulating immune function. The immune system has increasingly been recognized as a key player in early brain development^27,28^. In particular, the convergence of neurodevelopmental and immune function processes in brain regions with protracted developmental trajectories (e.g. prefrontal cortex)^29,30^ may help us contextualize our results of the genetic mechanisms contributing to the unique expansion of these brain regions. Indeed, we found genes involved in immunoregulation to be shared between surface area of the brain globally and dorsolateral prefrontal cortex regionally, such as the NF-κB pathway which responds to brain injury but also plays a role in brain plasticity and neurogenesis^31^. The role of the immune system, particularly microglia and other cytokines, in early prefrontal cortical development is thought to be important in shaping excitatory/glutamatergic neural circuits, where disruptions can confer later vulnerability to known neurodevelopmental disorders such as autism and schizophrenia^32^. We also found overlapping genes between global area and dorsolateral prefrontal cortex involved in GABAergic signaling, and for the growth factor receptor *EGFR*, which plays an important role in maintaining the progenitor pool in early brain development at a time when both neuronal and glial cells are being generated^33^. The functional profiling of these overlapping genes are consistent with the hypothesis that immune genes involved in prefrontal cortical development contribute to synaptic pruning and myelin growth events^34^.

Shared genes (e.g. *EIF4G1, PSMD2, RPSA, WNT3, WNT9B*) between global thickness and dorsolateral prefrontal thickness revealed an intriguing overlap in processes involved in Slit/Robo and WNT signaling, important for axon guidance and anterior-posterior patterning of the cortex, respectively^1,35^. Collectively these genes were also tied to molecular processes of dopaminergic differentiation, which provides an additional layer of evidence that may help contextualize our finding from gene ontology analyses suggesting enrichment for genes involved in tangential migration underlying dorsolateral prefrontal thickness.

Our results showcase how adjusting for global brain measures in GWAS models can change the interpretation of the biological pathways underlying regional brain morphology. Our gene ontology analyses exemplifies this, with common and unique biological processes associated with each region between the two GWAS approaches. Notably, the common processes are related to neurodevelopment. On the other hand, we find a large proportion of gene ontology terms associated with cancer and cell signaling pathways underlying many of our brain phenotypes in GWAS analyses that retain the global signal (e.g. do not adjust for global measures), which overlaps with the gene ontology terms found for global surface area (indexed by hashtags # in Fig. 3B). Several genes contributed to these cell signaling pathways, which are also observed in databases related to cancer, such as *FOXO3, PIK3CD, AKT3* and *TCF7L1*. These genes are relevant to neurodevelopment as they are involved in general cell cycle maintenance processes, such as *TCF7L1* for regulation of cell cycle genes^36^, and *PIK3CD* playing a role in signaling cascades involved in cell growth, survival and proliferation^37^.

Our findings also highlight the importance of studying inversion polymorphisms, particularly on 17q21.31, when mapping brain morphology. Consistent with Shin et al 2020’s GWAS findings of the first principal component explaining a large proportion of variance in surface area data7, we also found an enrichment of SNPs associated with global surface area located on chromosome 17. Global expansion of the brain and higher order cognitive regions in early development could be enriched for genes linked to the 17q21.31 inversion region. For instance, we see six shared genes in this inversion polymorphism region between global and dorsolateral prefrontal area (i.e. *SPPL2C, MAPT, STH, KANSL1, ARL17B, LRRC37A*), which together are involved in morphology of apical dendritic contacts, and in turn, neuronal communication. Four of these six genes show stronger effects (i.e. explain more variance) in dorsolateral prefrontal area compared to global area, supporting the idea that genes in the 17q21.31 inversion region may contribute to the disproportionate expansion of prefrontal cortical regions.

Genetic correlations between global measures and individual brain regions recapitulated cortical hierarchy and complemented our previous work^9–11^. We show once again an anterior/posterior gradient of genetic correlations with total surface area and regional area, alongside a dorsal/ventral pattern of correlations with mean cortical thickness and regional thickness. The latter thickness gradient has also been independently demonstrated in a twin study of cortical topological organization ^38^. Altogether our results suggest the important role of global measures in understanding this known cortical hierarchical structure.

We acknowledge several limitations in our current study. Firstly, we include several covariates in our models that do correlate with global brain measures, namely age and sex. It is well-established that males tend to have larger brain size, thus regressing out sex may contribute to removal of some global effects, even in our GWAS*g+r* models. Including age as a covariate may also have a similar effect, especially for cortical thickness, although the majority of our sample is within an age range that precedes steep declines in cortical thinning. Secondly, our included sample is of European descent, although we have previously shown in our work that GWAS results largely replicate in a sample of diverse ancestry. Thirdly, we acknowledge that our interpretation of results is impacted by our chosen gene mapping strategy and interactome database selection. The method we used, MAGMA, links genes based on their spatial proximity to a SNP, but may not always be the most relevant gene^39^; although there have been reports for specific traits where the likelihood of the nearest gene being causal is high^40^. Accurate gene mapping is a challenge in the field of genetics more broadly^41^, but we have included results from other gene mapping strategies in our supplementary material to offer readers resources from other tools, and found that these alternative mapping strategies provided similar interpretation of results.

Given that many brain imaging and genetics studies of structural brain features regress out global measures^42,43^, this study provides a more nuanced view of how global measures add to our interpretation of the genetics underlying brain morphology. The overlap in genetic architecture between global measures and multiple distributed brain regions could also be further elucidated with multivariate methods in future work, given its success in boosting discovery of novel genetic variants underlying cortical morphology^44–46^. Understanding global as compared to regional patterns of cortical morphology can have important implications for our understanding of higher cognitive function^47^ and in turn, psychiatric and neurological disorders that impact such cognitive processes.

## ONLINE METHODS

### Sample

Imaging and genomics data were taken from participants of the UK Biobank (UKB) population cohort, obtained from the data repository under accession number 27412^48–51^. All participants gave written informed consent^51^. The composition, set-up, and data gathering protocols of the UKB have been extensively described elsewhere^48–50^. For this study, White Europeans were selected that had undergone the neuroimaging protocol. We excluded 1094 individuals with a primary or secondary ICD10 diagnosis of a neurological or mental disorder, as well as 594 individuals with bad structural scan quality as indicated by an age and sex-adjusted Euler number^12^ more than three standard deviations lower than the scanner site mean. Our final sample included 32,488 individuals (mean age=64.2 [range: 45.1-81.8, SD: 7.5], %female=52.2).

### MRI processing and atlas definition

T1-weighted scans were collected from three scanning sites throughout the United Kingdom on identical Siemens Skyra 3T scanners with a 32-channel receive head coil. The UKB core neuroimaging team has published extensive information on the applied scanning protocols and procedures^50^. The T1 scans were obtained from the UKB data repositories and stored locally at the secure computing cluster of the University of Oslo. The standard “recon-all -all” processing pipeline of Freesurfer v5.3 was applied to perform automated surface-based morphometry segmentation^52^. Both surface area and cortical thickness are defined at the vertex-level, with surface area extracted from the white surface, and cortical thickness calculated as the distance between the white surface and pial surface.

Twelve regions of interest were extracted per hemisphere based on two genetically-informed atlases for cortical thickness and surface area for both discovery and replication cohorts. These atlases have been previously developed by our group, using a data-driven fuzzy clustering technique to identify parcels of the human cortex that are maximally genetically correlated based on the MRI scans of over 400 twins. No spatial information was used in creating the atlases. The only information used in deriving the parcels was the genetic correlations of cortical thickness or surface area among all vertices. More details can be found in our previous work^9,10^. We combined cortical phenotypes across hemispheres, as we have previously presented^11^.

Before running the GWAS on each measure, we regressed out age, sex, scanner site, a proxy of scan quality (FreeSurfer’s Euler number)^12^, and the first ten genetic principal components from each measure. We also regressed out whether or not the individual had a brain diagnosis based on ICD10 diagnostic information collected by the UKB (https://biobank.ndph.ox.ac.uk/showcase/field.cgi?id=41202). Individuals were classified as having a brain diagnosis if they met criteria for at least one class F (mental and behavioral disorders) or class G (disorders of the nervous system) diagnosis, with the exception of G56 -carpal tunnel syndrome, which is an extremely common condition and thus we did not consider it as a neurological diagnosis. Approximately 8% of our UKB sample were classified as having a mental/behavioral or neurological disorder. Subsequently, we applied a rank-based inverse normal transformation to the residuals of each measure, ensuring normally distributed input into each GWAS.

We denote one set of analyses as GWAS*r* for regional associations only after adjusting for global measures (i.e. total surface area, mean whole brain thickness) in the pre-residualized phenotypes, and the second set as GWAS*g+r*, which does not adjust for globals (Figure 1A).

### Contributions of global measures to model fits per phenotype

Four sets of tests were used to compare model fits when including and excluding global measures for our cortical phenotypes. These models all included the 15 covariates outlined above (age, sex, scanner, brain diagnosis, Euler number, 10 PCs). First, the Akaike Information Criterion (AIC) was applied^53^. AIC is often used by stepwise variable selection models and estimates the relative amount of information lost by a given model. We then applied the Bayesian information criterion (BIC)^54^, which is another criterion for comparing regression models. For both AIC and BIC, the smaller the value, the better the model. Next we used an ANOVA F test, which compares a full model and its reduced model, with the null hypothesis that the simpler (reduced) model is as good as the full model. Finally, we applied the Least Absolute Shrinkage and Selection Operator (LASSO)^55^, which uses cross validation to calculate the mean square error (MSE) for every combination of predictors, and then selects the best regression model with the minimum MSE. We first fixed all 15 covariates or predictors, to see if LASSO would select the global variable (mean thickness or total surface area) into the best model. Finally, this restriction was released to allow LASSO to select from all 16 predictors.

### Genotype quality control and imputation

The UKB v3 imputed data was used, which has undergone extensive quality control procedures as described by the UKB genetics team^51^. After converting the BGEN format to PLINK binary format, additional standard quality check procedures were carried out, including removal of single nucleotide polymorphisms (SNPs) with low imputation quality scores, filtering out individuals with more than 10% missingness, SNPs with more than 5% missingness, and SNPs failing the Hardy-Weinberg equilibrium test at p=1e-6. A minor allele frequency (MAF) threshold of 0.01 was used, leaving 7,853,566 SNPs. To further account for population substructure, we used results from UKB-specific principal components analysis (PCA), which were generated with flashPCA^56^.

### GWAS of cortical phenotypes for both GWAS*r* and GWAS*g+r*

We used fastGWA implemented in GCTA^57^, a mixed linear model-based tool, that controls for population stratification by principal components and takes into account relatedness using a sparse genetic relationship matrix (GRM). This method extends linear mixed model-based association analysis for use of biobank-scale data in a resource-efficient manner. The genome-wide significant loci were defined by clumping in PLINK^13^ (r^2^=0.1, distance=250kb) and thresholded at p<5e-8. Hits corrected for multiple comparisons can be found in our previous work^11^.

We then computed genetic correlations using LDSC^58^ per region between GWAS*g* and GWAS*g+r* (Figure 1B). We also computed genetic correlations per region between GWAS*r* and GWAS*g+r* summary statistics, where only the former set of GWAS regresses out the global signal (Figure 1C).

### Functional follow-up with FUMA

We used the web tool Functional Mapping and Annotation of Genome-Wide Association Studies (FUMA) (https://fuma.ctglab.nl/)^59^ to conduct a generalized gene-set analysis using MAGMA within FUMA^60^ to generate gene lists from GWAS*r* and GWAS*g+r* results, as well as for gene lists obtained for global area and thickness. For each gene, SNPs were selected that were within exonic, intronic and untranslated regions as well as SNPs within 50 kb upstream and downstream of the gene, a window size that has been used in previous cortical GWAS^14^. We used the 19,241 protein-coding genes in MAGMA in our main analysis. The gene-based p-value was calculated based on the mean of the summary statistic (*χ*^2^ statistic) of GWAS for the SNPs in a gene^60^. The *p-*value significance threshold was determined using the Bonferroni method, 0.05 divided by the number of genes (19,241), which is 2.6 × 10^−6^. The genes for GWAS*r* and the global measures have been previously reported in Makowski et al.

We also used FUMA to annotate significant SNPs from GWAS*r*, GWAS*g* and GWAS*g+r*, using positional, eQTL and chromatin interaction mapping. Summaries and details of these gene lists can be found in Supplementary Tables 16-18. Additional information on these three mapping strategies are as follows:

i. Positional mapping of SNPs, whereby SNPs within a 10kB window from known protein-coding genes in the human reference assembly (GRCh37/hg19) are mapped;
ii. eQTL mapping whereby allelic variations at a SNP is significantly linked to expression of a gene, where we considered eQTLs within cortical structures from GTEx v8, the UK Brain Expression Consortium (http://www.braineac.org/), the Common Mind Consortium ^61^, and PsychENCODE ^62^ (http://resource.psychencode.org);
iii. chromatin interaction mapping, to assess interactions between chromatin state within 200bp accessible for transcription and other regions of the 3D genome, using data from dorsolateral prefrontal cortex and neural progenitor cells (GSE87112), PsychENCODE, and adult and fetal cortex ^63^.

### Identifying overlap in gene modules between global and regional GWAS results

We quantified the genetic overlap between global area/thickness and each of the brain regions using the human protein interactome from STRING (a database of protein-protein interaction networks)^64^ and the genes we identified through MAGMA for each phenotype to create genetic modules for each brain trait. This network-based mapping approach can provide more biologically meaningful information compared to simply looking at overlapping genes between two traits. STRING consists of both physical and functional interactions, derived through co-expression, biological knowledge databases, and computational techniques. Interactions are scored based on accumulation of different types of evidence^64^. In our analysis we used interactions classified as ‘high confidence’ (combined score > 0.7), for the human interaction version 11.0, containing 17,185 proteins and 420,534 interactions^17^.

We assessed whether the genes underlying global measures were overlapping or separated from genes underlying regional brain measures (i.e. from GWAS*g+r*) by using a network separation analysis adapted from Menche and colleagues^17^. First, we calculated the mean shortest distance, dAB, between each global (A) and regional brain region (B) pair, as well as the shortest distance between genes within each of the brain phenotype modules (dAA and dBB). To take into account the size of each individual module, we calculated separation (SAB) between global and regional module pairs with the following formula: SAB=dAB -((dAA +dBB)/2). To assess whether global and regional genes demonstrated more significant overlap than expected by chance, 500 random gene sets with network degree-matched genes (i.e. genes with the same number of interactions) as the initial test set were generated to yield a distribution from which test statistics could be calculated from, similar to work from our group^25^ and others^17^. In addition to using gene lists generated by MAGMA, we also computed separation statistics using genes mapped by FUMA, which includes intergenic mapping (Supplementary Tables 16-18). Visualization of results was done through Cytoscape^19^ and functional profiling was completed with g:Profiler^18^ (Figure 1D).

## ACKNOWLEDGMENTS/DISCLOSURES

The authors would like to thank the research participants and staff involved in data collection of the UK Biobank. The authors would also like to thank Tobias Kaufmann for his help processing UK Biobank imaging data. This research was supported by the National Institutes of Health under R01MH118281, R01MH122688, R56AG061163. CM is supported by the Canadian Institutes of Health Research (CIHR) and the Kavli Institute for Brain and Mind (KIBM). PMV and JY acknowledge funding from the National Health and Medical Research Council (1113400) and the Australian Research Council (FT180100186 and FL180100072). JY is supported by the Westlake Education Foundation. DvdM is supported by Research Council of Norway (project #276082). **Author contributions:** Study design: C.M., H.W., D.vd.M., P.M.V., J.Y., C.H.C; Data analysis: C.M., H.W., Y. Q., D.vd.M., J.Z., C.H.C.; Manuscript writing: C.M., C.H.C.; UKB data processing: D.vd.M., O.F., Data visualization: C.M., A.S., A.Q., D.vd.M., C.H.C.; Manuscript revisions: all authors. **Data and materials availability:** UK Biobank data incorporated in this work was gathered from the public UK Biobank resource. All researchers who wish to access the research resource must register with UK Biobank by completing the registration form in the Access Management System (AMS) (https://www.ukbiobank.ac.uk/enable-your-research/register).

## Supplementary Tables

Makowski_etal_Supplementary_Tables_May2022.xlsx

## Notes

### Competing Interest Statement

The authors have declared no competing interest.

